# Impact of gestational low-protein intake on embryonic kidney microRNA expression and in the nephron progenitor cells of the male offspring fetus

**DOI:** 10.1101/2020.02.18.954800

**Authors:** Letícia de Barros Sene, Íscia Lopes-Cendes, Wellerson Rodrigo Scarano, Adriana Zapparoli, José Antônio Rocha Gontijo, Patrícia Aline Boer

## Abstract

Authors have demonstrated that gestational low-protein (LP) intake offspring showed a lower birth weight, reduced nephrons numbers and renal salt excretion, chronic renal failure, and arterial hypertension development when compared to normal (NP) protein intake group in adult life. The current study was aimed to evaluate the miRNAs and predicted gene expression patterns in the fetal kidney at 17 days of gestational (17-DG) protein-restricted male offspring (LP) in an attempt to elucidate the possible molecular pathways and disorders involved in renal cell proliferation and differentiation during kidney development. By NGS and RT-qPCR were identified, 44 differentially expressed miRNAs, which 19 miRNA were up- and 25 downregulated in LP compared to NP offspring metanephros. Among the top 10 deregulated miRNAs, the study selected 7 miRNAs which its biological targets were related to proliferation, differentiation, and cellular apoptosis processes. As could be seen below, both miRNA-Seq and TaqMan data analysis have shown a consistent change of miRNA expression in LP animals relative to control NP age-matched animals. By immunofluorescence, the LP fetus showed a significant 28% reduction of Six-2 labeled cells when compared to NP offspring associated with a reduced percentage of cell number and c-Myc metanephogenic cap and ureter bud immunostained cells in LP relative to NP offspring.

Additionally, the Ki-67 labeled area in CM was 48% lesser in LP compared to NP age-matched fetus. On the other hand, in LP, the CM and UB β-catenin marked area were 154 and 85% raised, respectively. Mtor immunoreactivity was also significantly higher in LP CM (139%) and UB (104%) compared to the NP fetus. In the LP offspring, the TGFβ-1 in UBs cells staining increased (about 30%). In contrast, Zeb1 metanephros-stained, located in the CM nuclei cells, enhanced 30% in LP and Zeb2 immunofluorescence, although present in whole metanephros structures, was similar in both experimental groups. In conclusion, the present study demonstrates that the miRNAs, mRNAs, and proteins are modified in LP animals with 17 DGs leading to the reduction of the reciprocal induction between CM and UB, and hence the number of nephrons. These findings will facilitate new functional approaches and further studies to elucidate the regulatory mechanisms involved in the processes of proliferation, differentiation, and apoptosis that occurs during renal development.

## Introduction

The lack of nutrients may result in pivotal pathways signaling changes during several stages of fetal development, which, in turn, may cause irreversible organ and system disorders in adulthood (Langley-Evans, 2006). Fetal Programming refers to any insult that occurs during a developmental period, and that causes long-term effects on the structure or function of an organism (Lucas, 1998). Disruptions in fetal programming result in low birth weight, fewer nephrons, increased risk of cardiovascular and renal disorders in adult age (Mesquita et al., 2010a,b; Sene et al.,2013, 2018). Other authors and us recently have demonstrated that gestational low-protein (LP) intake offspring showed a lower birth weight, 28% fewer nephrons, reduced renal salt excretion, chronic renal failure, and arterial hypertension development when compared to normal (NP) protein intake group in adult life (Schreuder et al., 2006; Zandi-Nejad et al., 2006; Mesquita et al., 2010a,b; Vaccari et al., 2013; Sene et al.,2013, 2018).

Nephrogenesis is a complex process involving tight control among genes expression, protein synthesis, and tissue remodeling process. The nephron number is established during renal embryonic development (Pan et al., 2017), being initiated by mutual interaction between the ureter bud (UB) and the metanephric mesenchyme (MM) progenitor cells (Grobstein, 1955; Saxén and Sariola, 1987). Signs from MM induce UB stimulated growth and a successive branch of tubule system. In turn, MM proliferation and differentiation constituting a mesenchymal cap (CM), is induced by UB ends (Phua et al., 2015a).

There is a critical interest in investigating the role of epigenetics phenomena concerning the long-term effects of prenatal stress on fetal development (Monk et al., 2012). MicroRNAs (miRNAs) are genomic-encoded small noncoding RNAs of approximately 22 nucleotides in length that play an essential role in post-transcriptional regulation of target gene expression (Bartel, 2004; Ambros, 2004, 2012; Bushati & Cohen, 2007). miRNAs control gene expression post-transcriptionally by regulating mRNA translation or stability in the cytoplasm (Pillai et al., 2007; Nilsen, 2007). Functional studies indicate that miRNAs are involved in critical biological processes during development and in cell physiology (Bushati & Cohen, 2007; Bartel, 2004), and changes in their expression are observed in several pathologies (Bushati & Cohen, 2007; Chang & Mendell, 2007).

Thus, miRNAs characterization has permitted gene regulation understanding, as well the process of cells proliferation, differentiation, and apoptosis, and explains pathophysiology disorder, including kidney disorders (Chu and Rana, 2007; Filipowicz et al., 2008; Kim et al., 2009; Li et al., 2010).

In the kidney ontogenic context, several researchers have demonstrated that miRNAs are indispensable for nephron development (Chu et al., 2014; Harvey et al., 2008; Lv et al., 2014; Marrone et al., 2014). Prior studies have shown that during kidney development, the underexpression of miRNAs in MM progenitor cells results in a premature reduction of cell proliferation and, consequently, decreased the number of nephrons (Ho et al., 2011; Nagalakshmi et al., 2011). This phenomenon is characterized by increased apoptosis and high expression of Bim protein in progenitor cells (Ho et al., 2011). Thus, we may suppose that miRNAs modulate the balance between apoptosis and the survival of these metanephric primary cells (Ho and Kreidberg, 2012).

Taking into account the above studies, unclear epigenetic phenomena and miRNA expression profiling are associated with kidney developmental disorder in maternal protein-restricted offspring. Thus, the current study was aimed to evaluate the miRNAs and predicted gene expression patterns in the fetal kidney at 17 days of gestational (17-DG) protein-restricted male offspring in an attempt to elucidate the possible molecular pathways and disorders involved in renal cell proliferation and differentiation during kidney development.

## Material and Methodology

### Animal and Diets

The experiments were conducted as described in detail previously (Lutaif et al., 2015) on an age-matched female and male rats of sibling-mated Wistar *HanUnib* rats (250–300 g) that were allowed free access to water and standard rodent chow (Nuvital, Curitiba, PR, Brazil). The Institutional Ethics Committee (#446-CEUA/UNESP) approved the experimental protocol, and the general guidelines established by the Brazilian College of Animal Experimentation were followed throughout the investigation. The environment and housing presented the right conditions for managing their health and well-being during the experimental procedure. Immediately after weaning at three weeks of age, animals were maintained under controlled temperature (25°C) and lighting conditions (07:00–19:00h) with free access to tap water and standard laboratory rodent chow (Purina rat chow: Na+ content: 135 ± 3µEq/g; K+ content: 293 ± 5µEq/g), for 12 weeks before breeding. It was designated day 1 of pregnancy as the day in which the vaginal smear exhibited sperm. Then, dams were maintained ad libitum throughout the entire pregnancy on an isocaloric rodent laboratory chow with either standard protein content [NP, n = 36] (17% protein) or low protein content [LP, n = 51] (6% protein). The NP and LP maternal food consumption were determined daily (subsequently normalized for body weight), and the bodyweight of dams was recorded weekly in both groups. Dams were euthanized in the 17 days of gestation (DG), and the fetus was removed, weighted and, tail and limbs were collected for sexing. The metanephros is collected for Next Generation Sequencing (NGS), RT-qPCR, and immunohistochemistry analyses.

### Sexing determination

The present study was performed only in male 17-DG offspring, and the sexing was determined by Sry conventional PCR (Polymerase Chain Reaction) sequence analysis (Miyajima et al., 2009). The DNA was extracted by enzymatic lysis with proteinase K and Phenol-Chloroform. For reaction, the Master Mix Colorless - Promega was used, with the cycling conditions indicated by the manufacturer. The Integrated DNA Technologies (IDT) synthesized the primer following sequences bellow:

> Forward: 5’-TACAGCCTGAGGACATATTA-3’
>
> Reverse: 5’-GCACTTTAACCCTTCGATTAG-3’.

### Total RNA extraction

RNA was extracted from NP (n = 4) and LP (n = 4) whole kidneys using Trizol reagent (Invitrogen), according to the instructions specified by the manufacturer. Total RNA quantity was determined by the absorbance at 260 nm using a nanoVue spectrophotometer (GE Healthcare, USA). RNA Integrity was ensured by obtaining an RNA Integrity Number - RIN > 8 with Agilent 2100 Bioanalyzer (Agilent Technologies, Germany).

### miRNA transcriptome sequencing (miRNA-Seq) and data analysis

Sequencing was performed on the MiSeq platform (Illumina). The protocol followed the manufacturer’s instructions available in (http://www.illumina.com/documents//products/datasheets/datasheet_truseq_sample_prep_kits.pdf). Briefly, the sequencing includes library construction, and this was used 1µg total RNA. In this step, the adapters are connected, the 3 ‘and 5’. After ligation of adapters, a reverse transcription reaction was performed to create cDNA, and it was then amplified by a standard PCR reaction, which uses primers containing a sequence index for sample identification — this cDNA library, subjected to agarose gel electrophoresis for miRNA isolation. After quantitation, the library concentration was normalized to 2 nM using 10 nM Tris-HCl, pH 8.5, and transcriptome sequencing were performed by MiSeq Reagent Kit v2 (50 cycles).

Data analysis was performed in collaboration with Tao Chen, Ph.D. from the Division of Genetic and Molecular Toxicological, National Center for Toxicological Research, Jefferson, AR, USA. The data from Next Generation Sequencing (NGS) of miRNAs were generated in FASTAQ format, and imported into BaseSpace.com (Illumina, USA). The data quality was evaluated using the base-calling CASAVA software developed by the manufacturer (Illumina). The analyzes were done by BaseSpace miRNA Analysis (from the University of Torino, Canada) and the sequence mapping of different miRNAs by Small RNA (Illumina, USA) for rat genome. The differentially expressed miRNA study was analyzed using Ingenuity Pathway Analysis software (Ingenuity, USA).

### miRNA expression validation

Four male offspring from different litters were used in each group for the miRNA (miR-127-3p, −144-3p, −298-5p, let-7a-5p, −181a-5p, −181c-3p, and −199a-5p) expression analysis. Total RNA was extracted from 17-DG samples using Trizol reagent (Life Technologies, USA), according to the instructions previously described (Chomczynski & Sacchi, 2006). Total RNA was quantified (Take 3 micro-volume plate of the Epoch spectrophotometer; BioTek®, USA). The RNA integrity was evaluated by electrophoresis on a denaturing agarose gel stained with GelRed Nucleic Acid Gel Stain (Uniscience, USA) and the RNA purity was assessed by the ratio of absorbance at 260 and 280 nm. Briefly, 450 ng RNA was reverse transcribed, without pre-amplification, using TaqMan® MicroRNA Reverse Transcription Kit and Megaplex RT Primers Rodent Pool A (Life Technologies, USA), according to the manufacturer’s guidelines. Complementary DNA (cDNA) was amplified using a TaqMan® Rodent MicroRNA Array A v2.0 with TaqMan Universal PCR Master Mix on QuantStudio 12K Flex System (Life Technologies, USA), according to the manufacturer’s instructions. Data analysis was performed using relative gene expression evaluated using the comparative quantification method (Livak & Schmittgen, 2001). The U87 gene was used as a reference gene. Mean relative quantity was calculated, and miRNAs differentially expressed between groups (LP-12d versus NP-12d and LP-16w versus NP-16w) were evaluated. miRNA data have been generated following the MIQE guidelines (Bustin et al., 2009).

### RT-qPCR of predicted target genes

Total RNA was extracted from 17-DG offspring kidneys using the Trizol method previously described (Chomczynski & Sacchi, 2006). The total RNA quantity, purity, and integrity were assessed as previously described for miRNAs expression analysis. For the cDNA synthesis, a High Capacity cDNA reverse transcription kit (Life Technologies, USA) was used. To analyze the level of gene expression Bax, Bim, Caspase-3, Collagen 1, GDNF, PCNA, TGFβ-1, Bcl-2, Bcl-6, c-myc, c-ret, cyclin A, Map2k2, PRDM1, Six-2, Ki67, MTOR, β-catenin, ZEB1, ZEB2, NOTCH1 and IGF1, the reaction of RT-qPCR was performed with SYBR Green Master Mix (Life Technologies, USA), using primers specific for each gene (Table 1). The reactions were done in a total volume of 20 µL using two µL of cDNA (diluted 1:30), ten µL SYBER Green Master Mix (Life Technologies, USA) and four µL of each specific primer (5 nM) StepOnePlus real-time PCR system (Applied BiosystemsTM). For real-time PCR, 2 µl cDNA (40ng/ µl) was added to a master mix comprising 10 µl TaqMan® Fast Advanced Master Mix (Life Technologies, EUA), 1 µl primer mix and 7 µl water for a reaction. Water was used in place of cDNA as a non-template control. The cycling conditions were: 50°C for 2 minutes, 95°C for 20 seconds, 50 cycles of 95°C for 1 second, and 60°C for 20 seconds. Amplification and detection were performed using the StepOne Plus (Life Technologies, EUA) and data acquired using the StepOne Software v2.1 (Life Technologies, EUA). Ct values were converted to relative expression values using the ΔΔCt method with offspring heart data normalized to GAPDH as a reference gene. IDT® Integrated DNA Technologies provided primers for mRNA RT-qPCR. Bioinformatics methods played a central role in miRNA target prediction (Krek et al., 2005; John et al., 2006). Numerous target prediction algorithms exploiting different approaches have been recently developed for the prediction of miRNA-mRNA interactions (Witkos et al., 2011). In the present study, *in silico* target prediction was performed for differentially expressed miRNAs using the combined analysis of three algorithms based on conservation criteria TargetScan (Lewis et al., 2005), microRNA.org (John et al., 2004) and PicTar (Krek et al., 2005; Lall et al., 2006). Results were taken from each search analysis and cross-referenced across all the three research results. To exclude the hypertension effect on gene expression, only targets predicted in both 12-day and 16-week old animals were considered. Furthermore, only targets genes expressed in cardiac tissue were used for the analysis. To offer experimental support to *in silico* predicted targets, we evaluated the gene expression by RT-qPCR and quantified the protein levels by western blot analysis.

**Table 1:** Sequence of the primers used for RT-qPCR, designed by the company IDT.

| **Antibody** | **Dilution** | **Company** |
| --- | --- | --- |
| **Anti-Six2 (11562-1-AP)** | 1:50 | Proteintech |
| **Anti-c-myc (NBP1-19671)** | 1:150 | Novus Biologicals |
| **Anti-Ki67 (ab16667)** | 1:100 | Abcam |
| **Anti-Bcl2 (ab7973)** | 1:100 | Abcam |
| **Anti-TGFβ-1 (sc-146)** | 1:50 | Santa Cruz |
| **Anti-B-catenina (ab32572)** | 1:500 | Abcam |
| **Anti-Zeb1 (sc-10572)** | 1:50 | Santa Cruz |
| **Anti-Zeb2 (sc-48789)** | 1:50 | Santa Cruz |
| **Anti-VEGF (NB100-664)** | 1:50 | Novus Biologicals |
| **Anti-Caspase-3 clivada (9664)** | 1:200 | Cell Signaling |
| **Anti-Ciclina A (sc-31085)** | 1:50 | Santa Cruz |
| **Anti-WT1 (sc-192)** | 1:50 | Santa Cruz |
| **Anti-mTOR** |  |  |

### Immunohistochemistry

After euthanasia, the bodies of the fetus (n =4 per group) were removed and placed in the fixative (4% paraformaldehyde in 0.1 M phosphate buffer pH 7.4) for 4h. After were washed in running water, and followed by 70% alcohol until processed. The materials were dehydrated, diaphanized, and included in paraplast. The paraplast blocks were cut into 5-µm-thickness sections. Histological sections were deparaffinized and processed for immunofluorescence and immunoperoxidase. For immunofluorescence, the sections were incubated with a blocking solution (8% fetal bovine serum, 2.5% bovine albumin, and 2% skimmed milk powder in PBS). Subsequently, incubated with the primary antibody (anti-Six2) diluted in PBS containing 1% skim milk overnight under refrigeration. After washing with PBS, the sections were incubated with specific secondary antibody, conjugated to the Alexa 488 fluorophore, diluted in the same buffer, containing 1% milk for 2 hours at room temperature. After successive washes with PBS, the slides were mounted with coverslips using the Vectashield fluorescent assembly medium (Vector Laboratories, Inc. Burlingame), and the fluorescence in the specimen was detected by laser confocal microscopy. The images were obtained using the Focus Imagecorder Plus system. For the c-myc, ki67, Bcl2, TGFβ-1, β-catenin, Zeb1, Zeb2, Caspase 3 cleaved, cyclin A and WT1 proteins, immunohistochemistry was performed. The slides were hydrated, and after being washed in PBS pH 7.2 for 5 minutes, the antigenic recovery was made with citrate buffer pH 6.0, for 25 minutes in the pressure cooker. The slides were washed in PBS. Subsequently, endogenous peroxidase blockade with hydrogen peroxide and methanol was performed for 10 minutes in the dark. The sections were rewashed in PBS. Blocking of non-specific binding was then followed, and the slides were incubated with a blocking solution (5% skimmed milk powder, in PBS) for 1 hour. The sections were incubated with the primary antibody (Table 2) diluted in 1% BSA overnight in the refrigerator. After washing with PBS, the sections were exposed to the specific secondary antibody, diluted in 1% BSA, for 2 hours at room temperature. The slides were washed with PBS. The cuts were revealed with DAB (3,3’-diaminobenzidine tetrahydrochloride, Sigma - Aldrich CO®, USA), after successive washing with running water, the slides were counterstained with hematoxylin, dehydrated and mounted with a coverslip, using Entellan®. The images were obtained using the photomicroscope (Olympus BX51) or a Zeiss LSM 780-NLO confocal on an Axio Observer Z.1 microscope (Carl Zeiss AG, Germany) from National Institute of Science and Technology on Photonics Applied to Cell Biology (INFABIC) at the State University of Campinas.

**Table 2.**
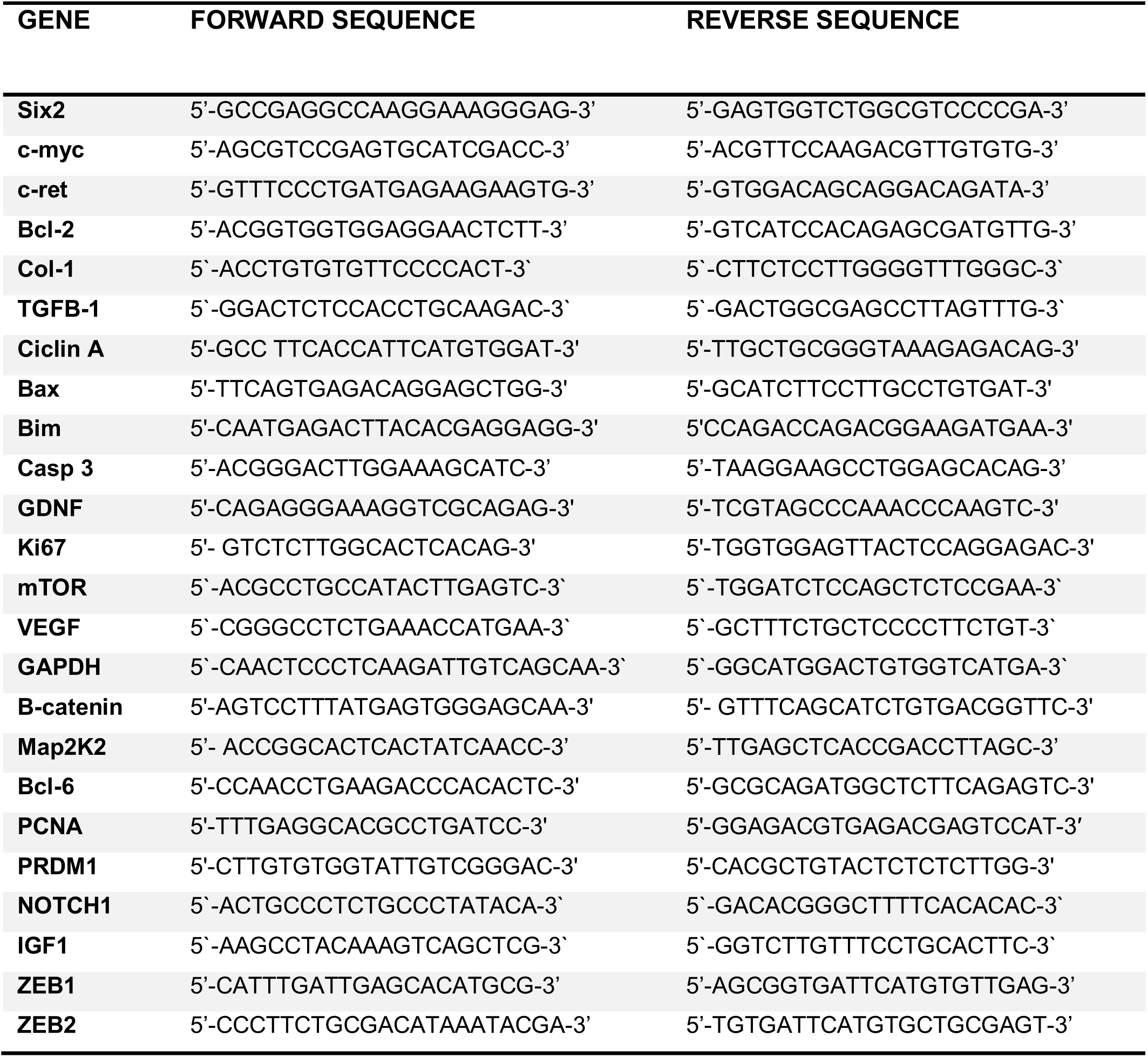
Dilution of antibodies used in immunohistochemistry.

### Morphology Quantification

Paraffin sections (5 µm) of the metanephros and nephrogenic area (5NP and 5LP from different mothers) as well CM and UB protein and cell quantifications, hematoxylin-eosin stained, were analyzed and microscopic fields digitized using CellSens Dimension software from a photomicroscope (Olympus BX51). We quantified all CM and UB of each metanephros analyzed (4NP and 4LP from different mothers), and statistical analysis was performed by t-test, and the values were expressed as mean ± SD. The p≤0.05 was considered significant. GraphPad Prism v01 Software, Inc., USA, was used for statistical analysis and graph construction.

### Statistical Analysis

Data are expressed as the mean ± standard deviation or as the median [lower quartile - upper quartile] and was previously tested for normality (Kolmogorov-Smirnov test) and equality of variance. Comparisons between two groups were performed using Student’s t-test when data were normally distributed, and the Mann-Whitney test when distributions were non-normal. Comparisons between two groups through the weeks were performed using 2-way ANOVA for repeated measurements test, in which the first factor was the protein content in the pregnant dam’s diet, and the second factor was time. When an interaction was found to be significant, the mean values were compared using Tukeýs post hoc analysis. Significant differences in miRNA expression were detected using a moderated t-test. Data analysis was performed with Sigma Plot v12.0 (SPSS Inc., Chicago, IL, USA). The significance level was 5%.

## Results

### Expression of miRNAs by miRNA-Sequencing (miRNA-Seq)

To understand the microRNA changes associated with maternal low-protein renal programming, we performed the expression of a global miRNA profiling analysis. It was identified 44 deregulated miRNAs (p ≤ 0.05), of which 19 and 25 miRNA, respectively, were up- or down-regulated (Table 3). The top expressed miRNAs, as well as its functions, pathways, and networks, were identified using Ingenuity Software (Table 4).

**Table 3.**
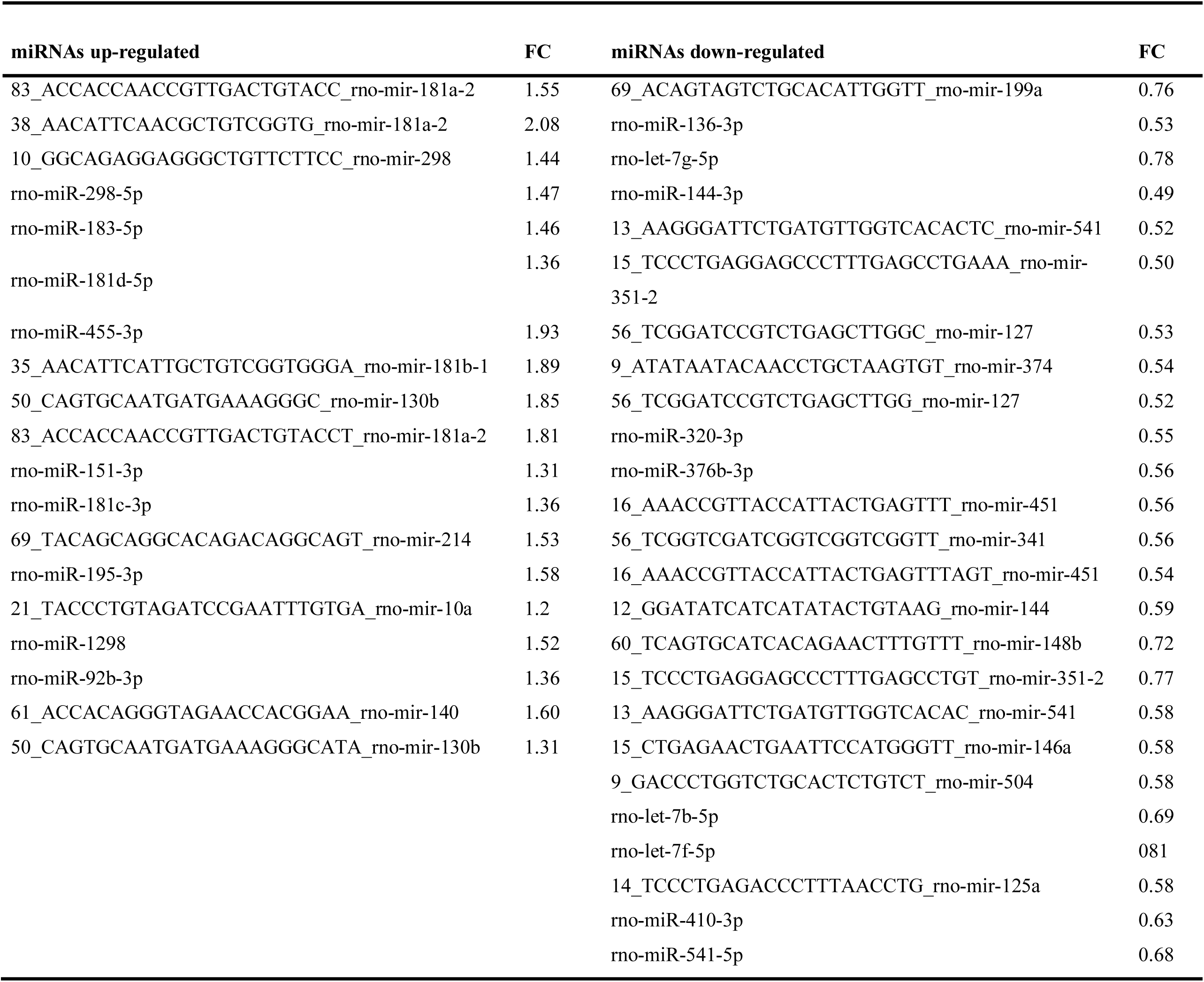
Lists of the deregulated miRNAs obtained by miRNA-Seq.

**Table 4.**
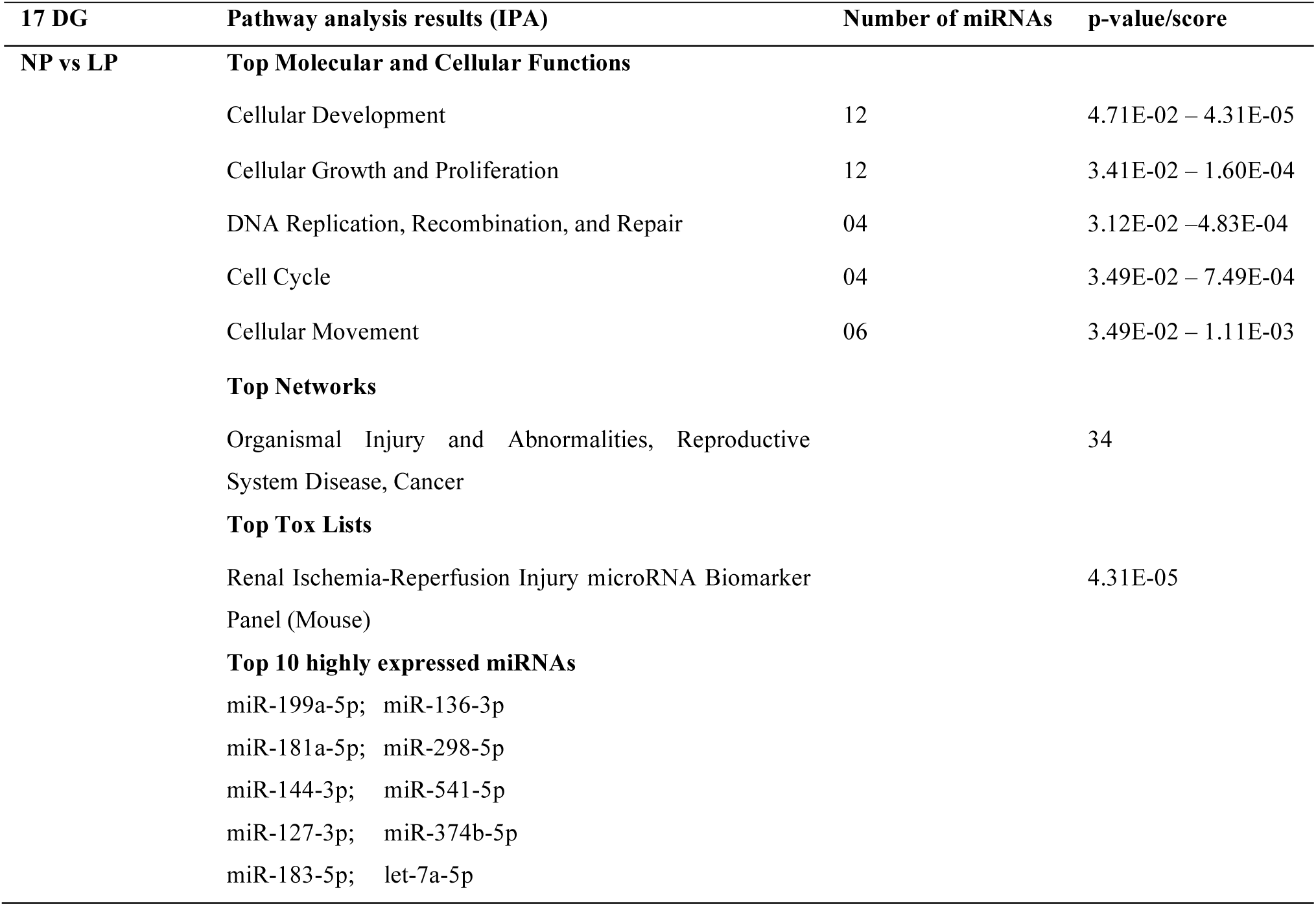
Top canonical pathways affected by differentially expressed miRNAs in 17 DG LP metanephros.

| 17 DG | Pathway analysis results (IPA) | Number of miRNAs | p-value/score |
| --- | --- | --- | --- |
| NP vs LP | **Top Molecular and Cellular Functions** |  |  |
|  | Cellular Development | 12 | 4.71E-02 – 4.31E-05 |
|  | Cellular Growth and Proliferation | 12 | 3.41E-02 – 1.60E-04 |
|  | DNA Replication, Recombination, and Repair | 04 | 3.12E-02 – 4.83E-04 |
|  | Cell Cycle | 04 | 3.49E-02 – 7.49E-04 |
|  | Cellular Movement | 06 | 3.49E-02 – 1.11E-03 |
|  | **Top Networks** |  |  |
|  | Organismal Injury and Abnormalities, Reproductive System Disease, Cancer |  | 34 |
|  | **Top Tox Lists** |  |  |
|  | Renal Ischemia-Reperfusion Injury microRNA Biomarker Panel (Mouse) |  | 4.31E-05 |
|  | **Top 10 highly expressed miRNAs** |  |  |
|  | miR-199a-5p; miR-136-3p |  |  |
|  | miR-181a-5p; miR-298-5p |  |  |
|  | miR-144-3p; miR-541-5p |  |  |
|  | miR-127-3p; miR-374b-5p |  |  |
|  | miR-183-5p; let-7a-5p |  |  |

### Validation of miRNA expression

*In* the animals of the LP group, let-7a-5p, miR-181a-5p, miR-181c-3p were upregulated, while the miR-127- 3p, miR-144-3p, and miR-199a-5p were downregulated relative to age-matched NP animals. The results do not show any difference in miR-298 expression, comparing both groups (Figure 1). Table 5 revealed the contrast of the values obtained by miRNAs sequencing with the RT-qPCR validation data. Although significant miRNA expression difference was observed in LP relative to NP offspring, the fold change (FC) of the validated miRNAs was similar to both techniques.

**Figure 1:**
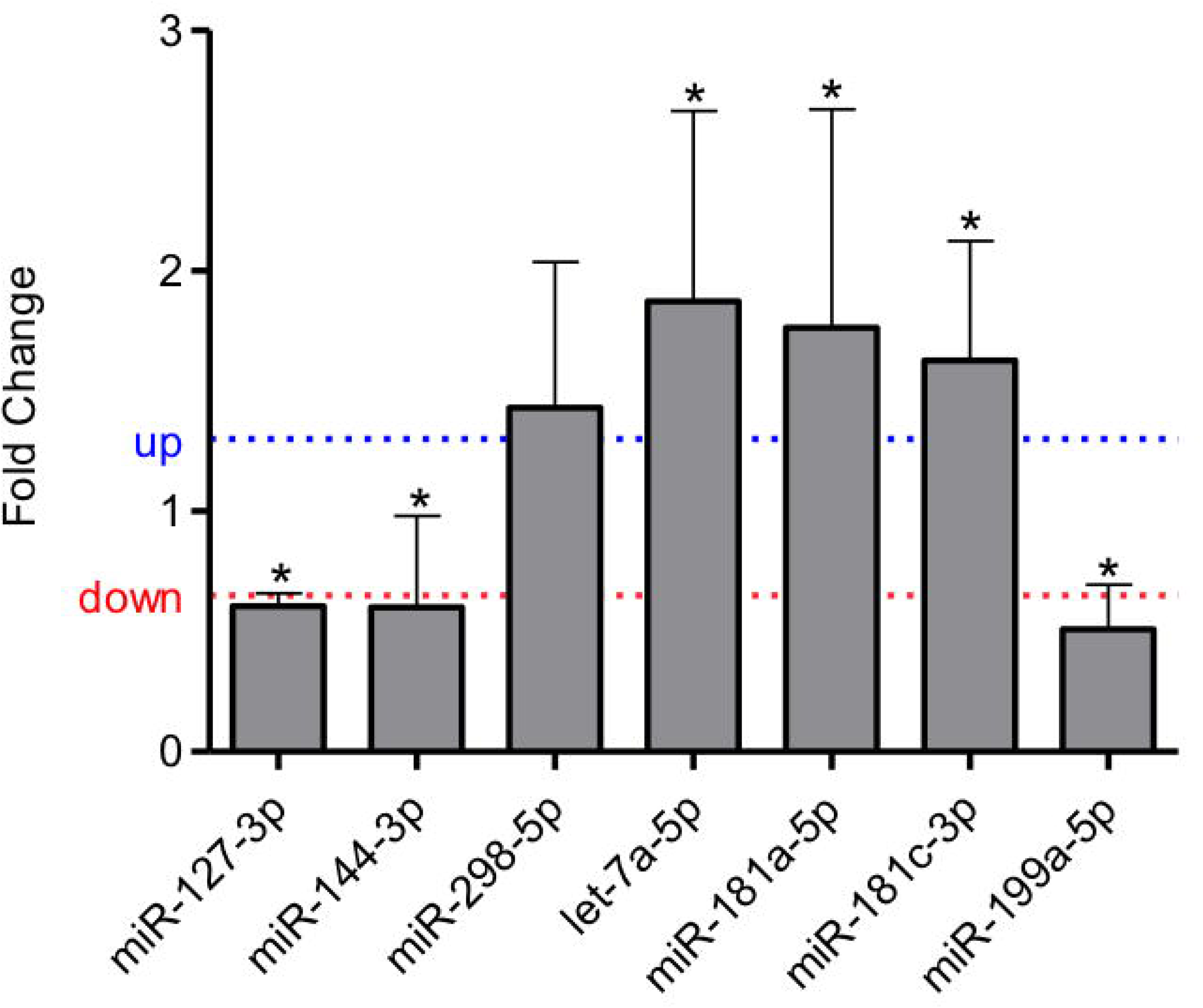
Expression of miRNAs in the metanephros from the 17th day LP fetus compared to their level of expression in the control group. The expression of each miRNA was normalized with U6 and U87. Data are expressed as fold change (mean ± SD, n = 4) concerning the control group. * p≤0.05: statistical significance versus NP.

**Table 5.**
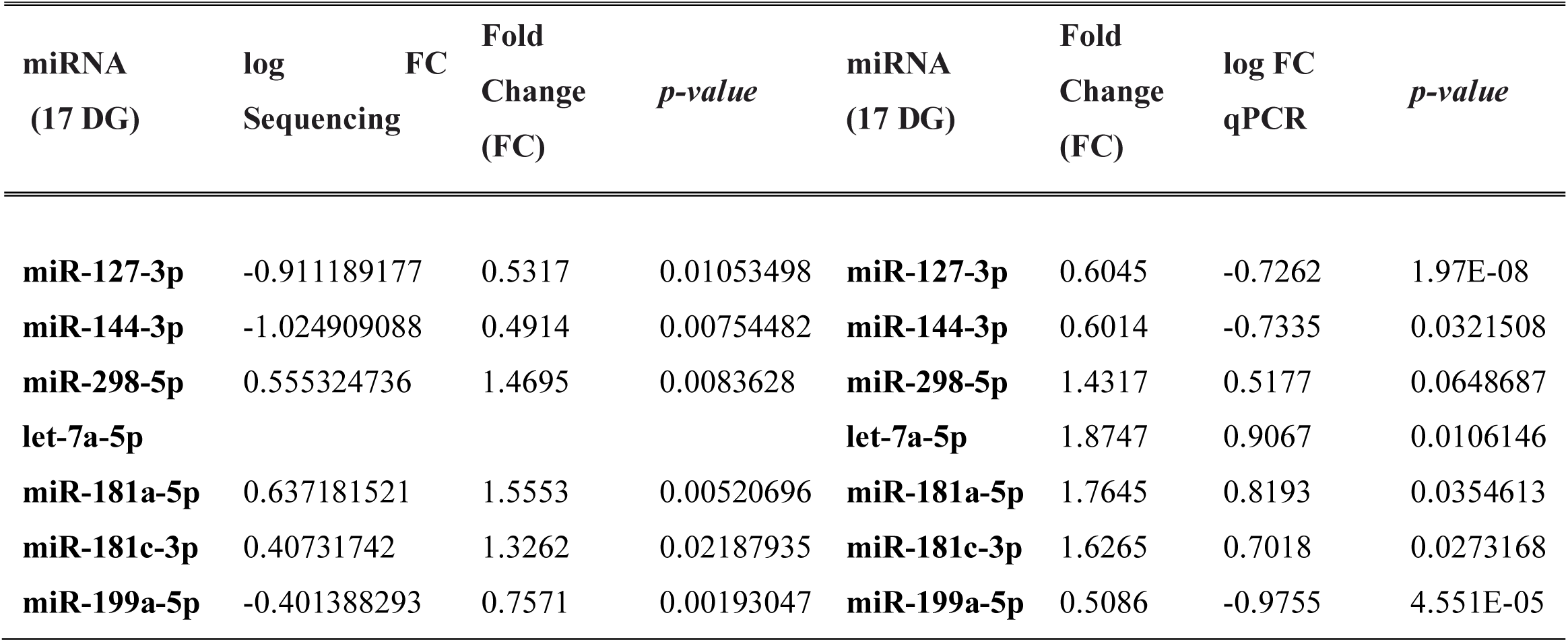
Comparison between the values obtained in the miRNA sequencing and the validation by RT-qPCR.

| **miRNA**<br>**(17 DG)** | **log**<br>**FC**<br>**Sequencing** | **Fold**<br>**Change**<br>**(FC)** | ***p-value*** | **miRNA**<br>**(17 DG)** | **Fold**<br>**Change**<br>**(FC)** | **log FC**<br>**qPCR** | ***p-value*** |
| --- | --- | --- | --- | --- | --- | --- | --- |
| **miR-127-3p** | -0.911189177 | 0.5317 | 0.01053498 | **miR-127-3p** | 0.6045 | -0.7262 | 1.97E-08 |
| **miR-144-3p** | -1.024909088 | 0.4914 | 0.00754482 | **miR-144-3p** | 0.6014 | -0.7335 | 0.0321508 |
| **miR-298-5p** | 0.555324736 | 1.4695 | 0.0083628 | **miR-298-5p** | 1.4317 | 0.5177 | 0.0648687 |
| **let-7a-5p** |  |  |  | **let-7a-5p** | 1.8747 | 0.9067 | 0.0106146 |
| **miR-181a-5p** | 0.637181521 | 1.5553 | 0.00520696 | **miR-181a-5p** | 1.7645 | 0.8193 | 0.0354613 |
| **miR-181c-3p** | 0.40731742 | 1.3262 | 0.02187935 | **miR-181c-3p** | 1.6265 | 0.7018 | 0.0273168 |
| **miR-199a-5p** | -0.401388293 | 0.7571 | 0.00193047 | **miR-199a-5p** | 0.5086 | -0.9755 | 4.551E-05 |

### miRNA-gene targets

The expressions of predicted targets of differentially expressed miRNA such as Six-2, Bcl-2, PRDM1, cyclin A, PCNA, GDNF, Collagen 1, Caspase 3 and Bim gene-expression level in LP fetus did not differ significantly from NP offspring. However, Bax, TGFβ-Bcl-6, c-ret, Map2k2, Ki67, mTOR, β-catenin, ZEB1, ZEB2, and IGF1 genes expression were upregulated in the LP group compared to controls. At the same time, c-myc and NOTHC1 were downregulated (Figure 2) in maternal protein-restricted offspring.

**Figure 2:**
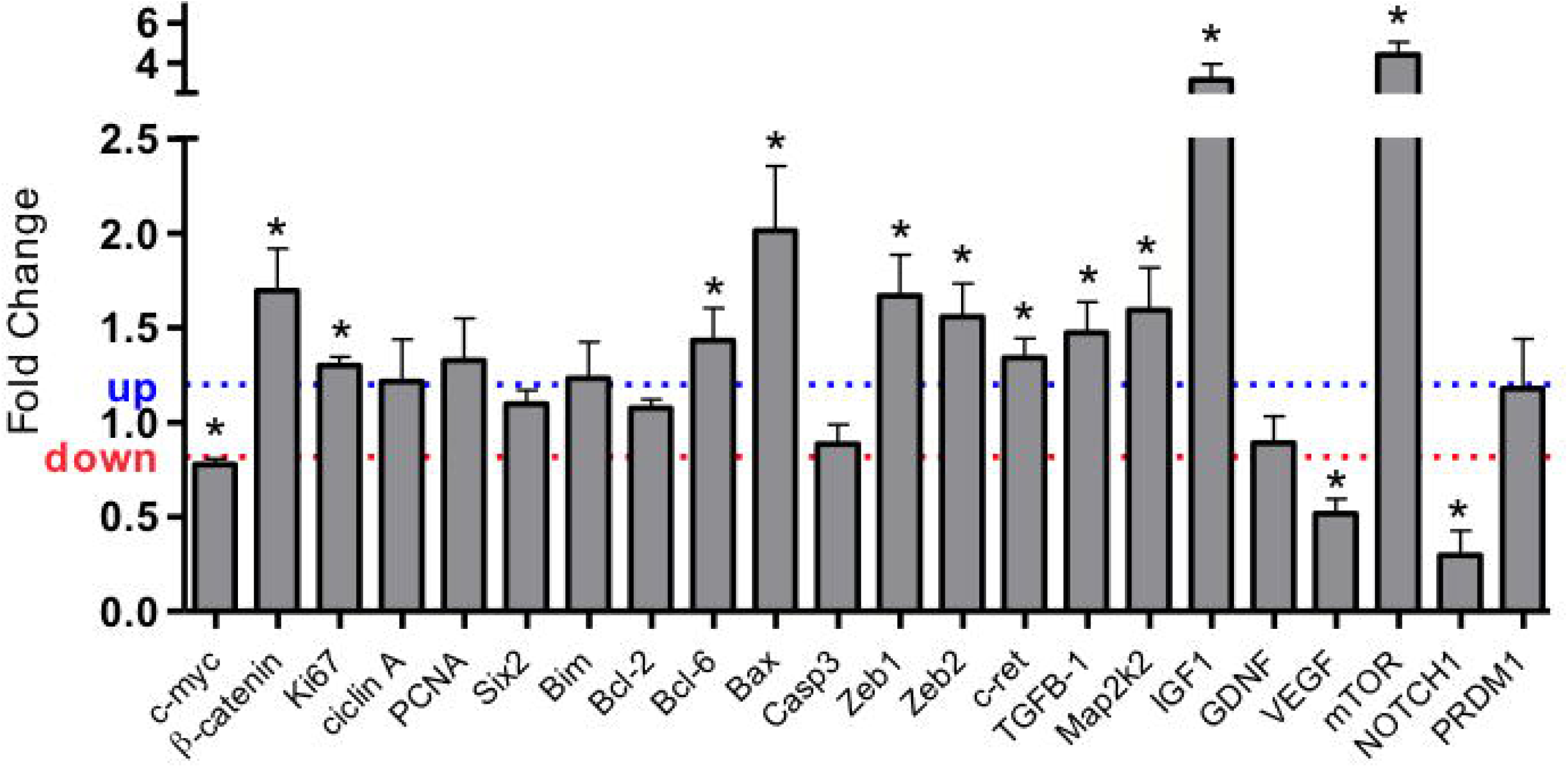
Expression of mRNA estimated by SyBR green RT-qPCR of metanephros from the 17th day LP fetus. The expression was normalized with GAPDH. Data are expressed as fold change (mean ± SD, n = 4) concerning the control group. * p≤0.05: statistical significance versus NP.

### Fetus body mass and metanephros morphometry

In the 17th DG, the offspring of the LP group presented no difference in body mass concerning to NP fetus. However, the LP fetus’s metanephros has shown a significant reduction (7.6%) in the nephrogenic area and the nephrogenic cortex thickness (about 29%) when compared to the NP group (Figure 3).

**Figure 3:**
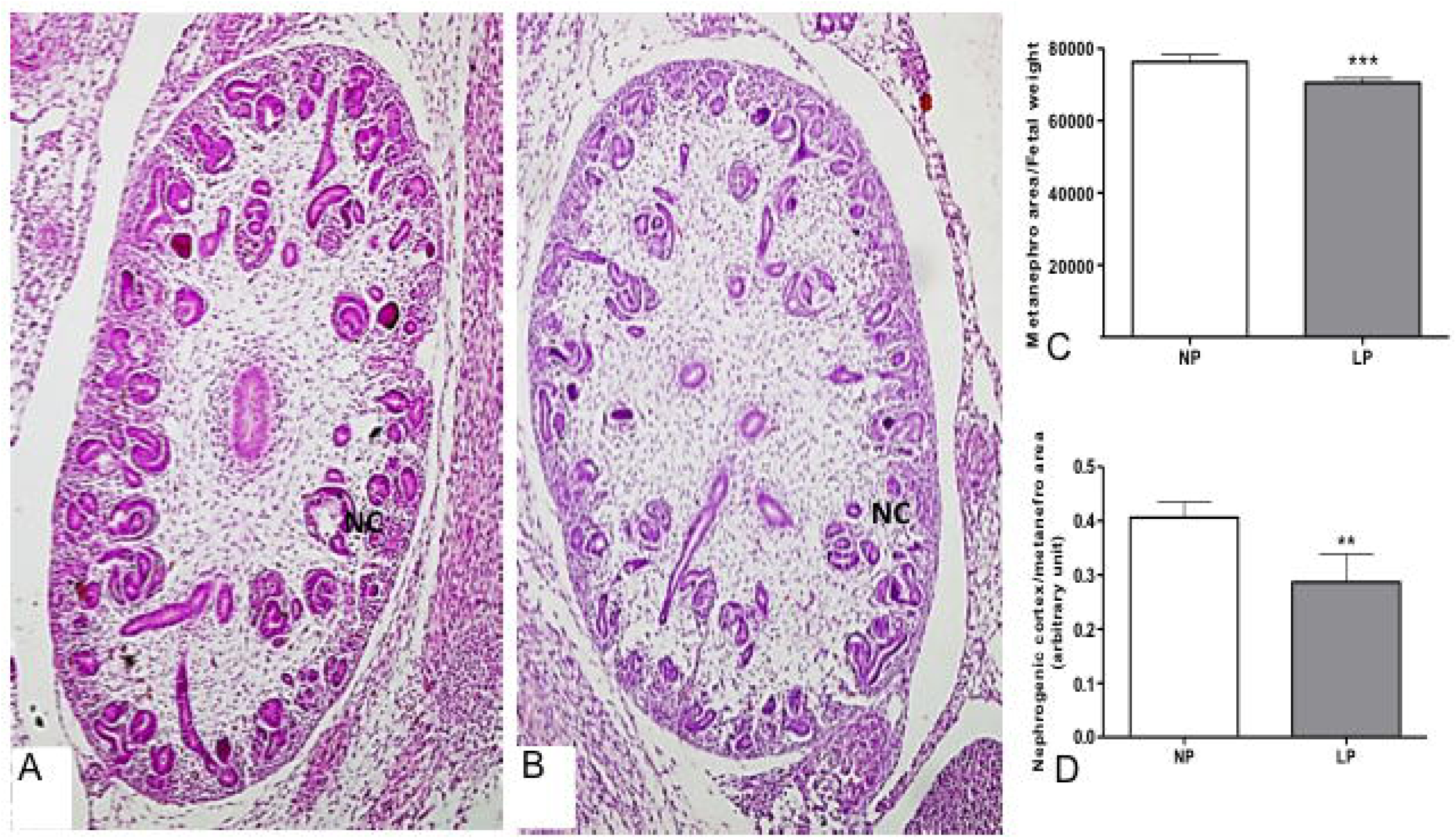
Metanephros of the fetus with 17 DG and quantifications. Comparing the HE stained micrography, we can observe the difference in the NP (A) and LP(B) metanephros size and nephrogenic cortex (NC) thickness. The differences between these parameters are statistically significant (C and D). **p<0.005; ***p<0.0001.

### Immunohistochemistry

By the immunofluorescence intensity quantification, the LP fetus showed a significant reduction (about 69%) of Six-2 labeled metanephros when compared to Six-2 metanephros-stained in NP offspring (Figure 4). Also, the Six-2 immunoperoxidase quantifications showed a reduced percentage of cell number (14%) and in metanephros from LP fetuses associated with 28% reduced Six2+ cells relative to NP offspring (Figure 4). Also, the present study showed a significant percent reduction of c-Myc CM and UB immunostained cells (less 14%) in LP relative to NP offspring (Figure 4). Additionally, the percentage of Ki-67 labeled area in CM was 48% lesser in LP compared to NP fetus, while Bcl-2 and cleaved caspase-3 immunoreactivity were not different from both groups (Figure 5-6). On the other hand, in LP, the CM and UB β-catenin labeled area were 154 and 85% raised, respectively, when compared to that availed in NP offspring (Figure 7). At the same time, mTOR immunoreactivity distribution also occupied a significantly more extensive area in LP CM (139%) and UB (104%) than in the NP fetus (Figure 7). In the LP offspring, the TGFβ-1 in UBs cells staining increased (about 30%), while in the CM, the immunostained cells were not different related to the NP group (Figure 8). The Zeb1 metanephros-stained, located in the CM nuclei cells, enhanced 30% in LP compared to the NP fetus (Figure 8). At the same time, the Zeb2 immunofluorescence, although present in whole metanephros structures, was similar in both experimental groups (Figure 8). In the current study, taking into account miRNA and mRNA expression and proteins immunostaining results present above may permit schedule representative pathway interactions to explain the observed findings (figure 9).

**Figure 4.**
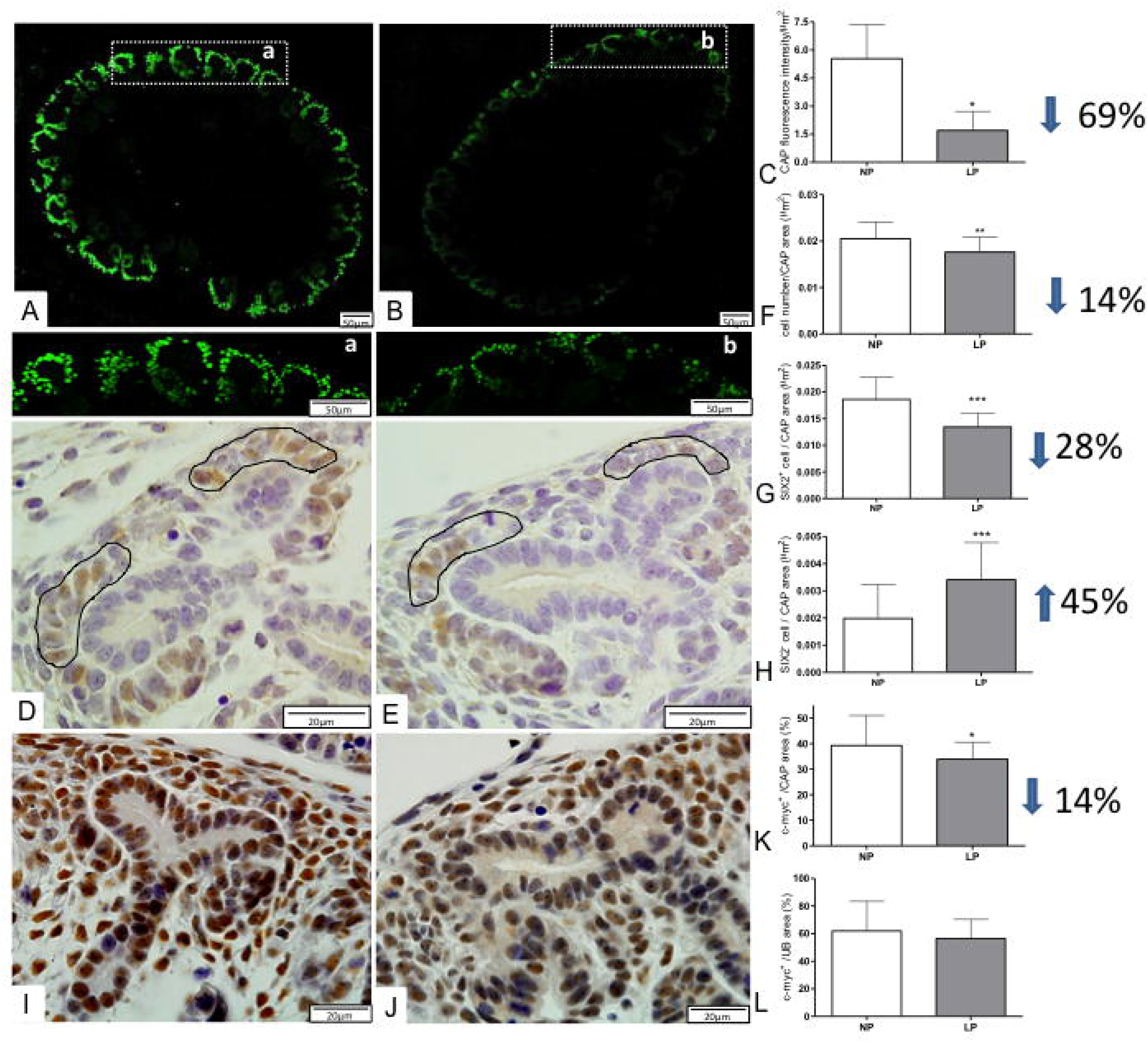
Immunofluorescence and immunoperoxidase for Six2 and c-Myc in metanephros of 17DG fetus. The Six2 immunofluorescence in LP (A, a) is reduced when compared to that observed in NP(B,b) metanephros and, when quantified, was significantly lower (C) in LP comparatively to NP. Additionally, after immunoperoxidase in NP (D) and LP (E) metanephros, we have counted the number of cells as well Six2 + cells in the caps (circled by black lines) that are significantly reduced in LP (F, G). In LP (J) c-Myc labeled area was reduced in the cap (K) but was the same in UB(L) when compared to NP (J). *p<0.005;**p<0.001; ***p<0.0001.

**Figure 5.**
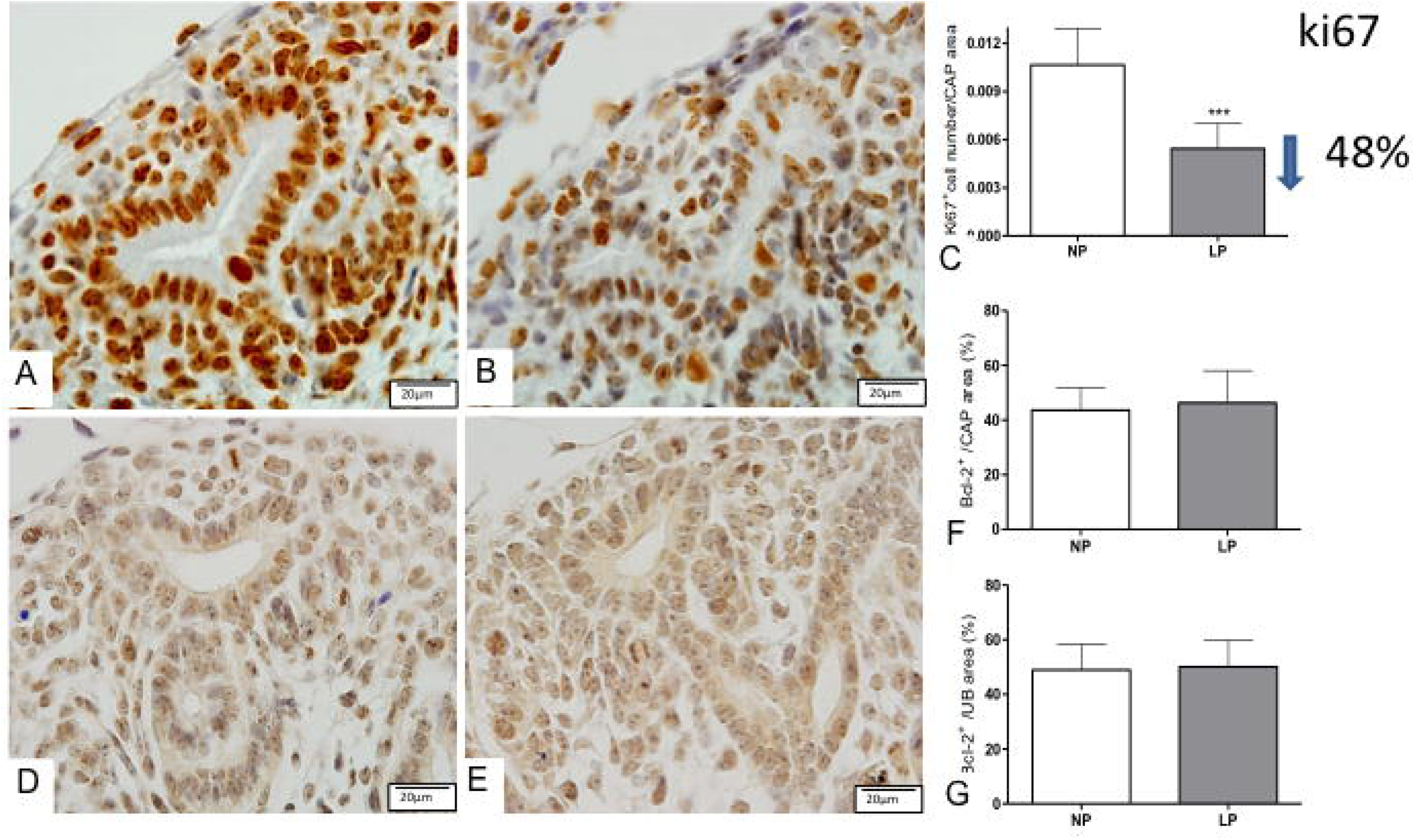
Immunoperoxidase for Ki67 and Bcl-2 in metanephros of 17DG fetus. The number of Ki67+ cells was reduced in LP (B) when compared to NP (A) and was statistically significant is the caps (C). The Bcl-2 labeled area was the same in NP (D) and LP (E) in both UB and cap (F, G). ***p<0.0001.

**Figure 6.**
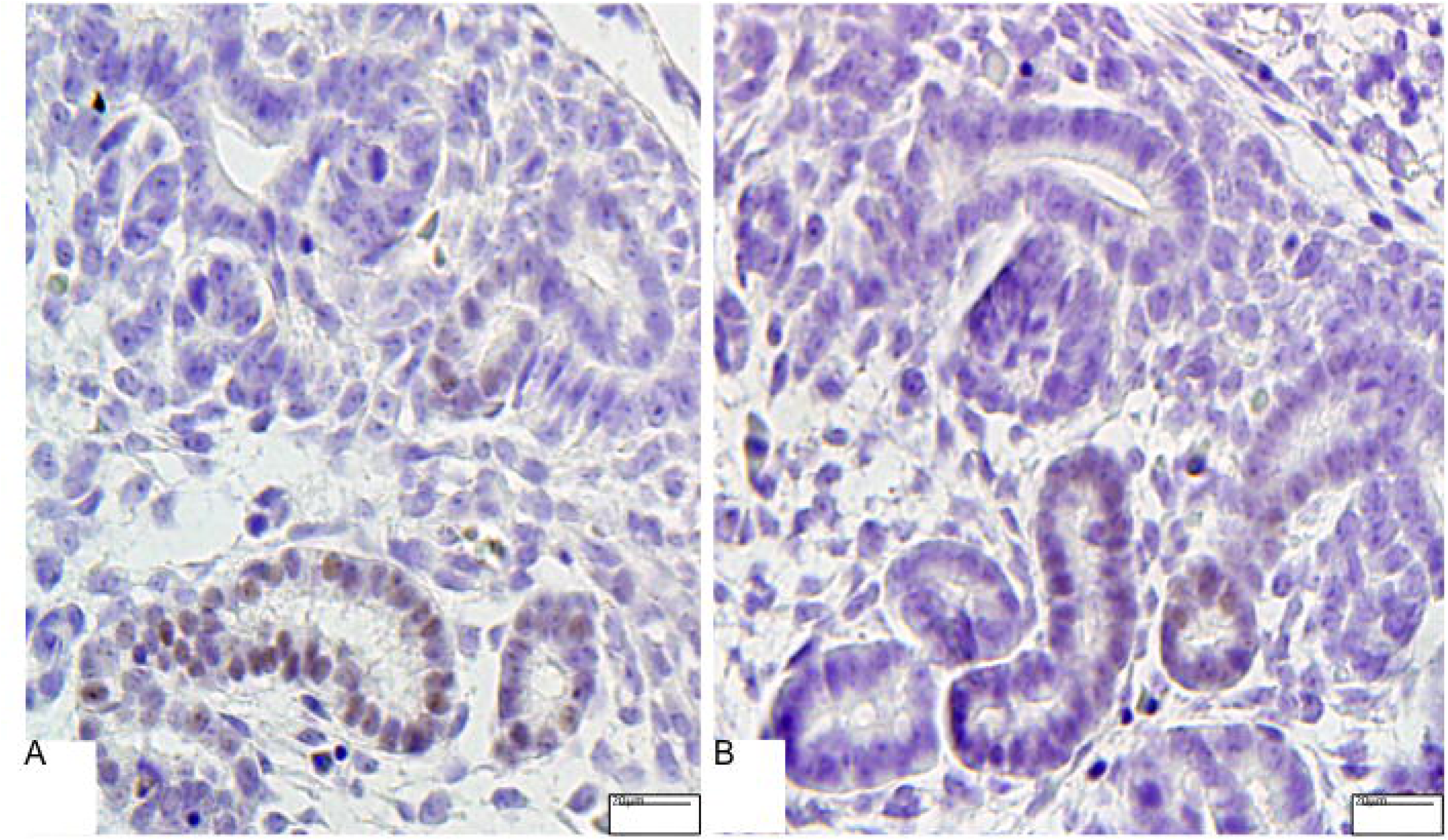
Immunoperoxidase for cleaved caspase 3 in metanephros of 17DG fetus. The positive cells were preferentially located in the ureteric trunk epithelium, and apparently, the quantity is not different in LP (B) when compared to that observed in NP (A).

**Figure 7.**
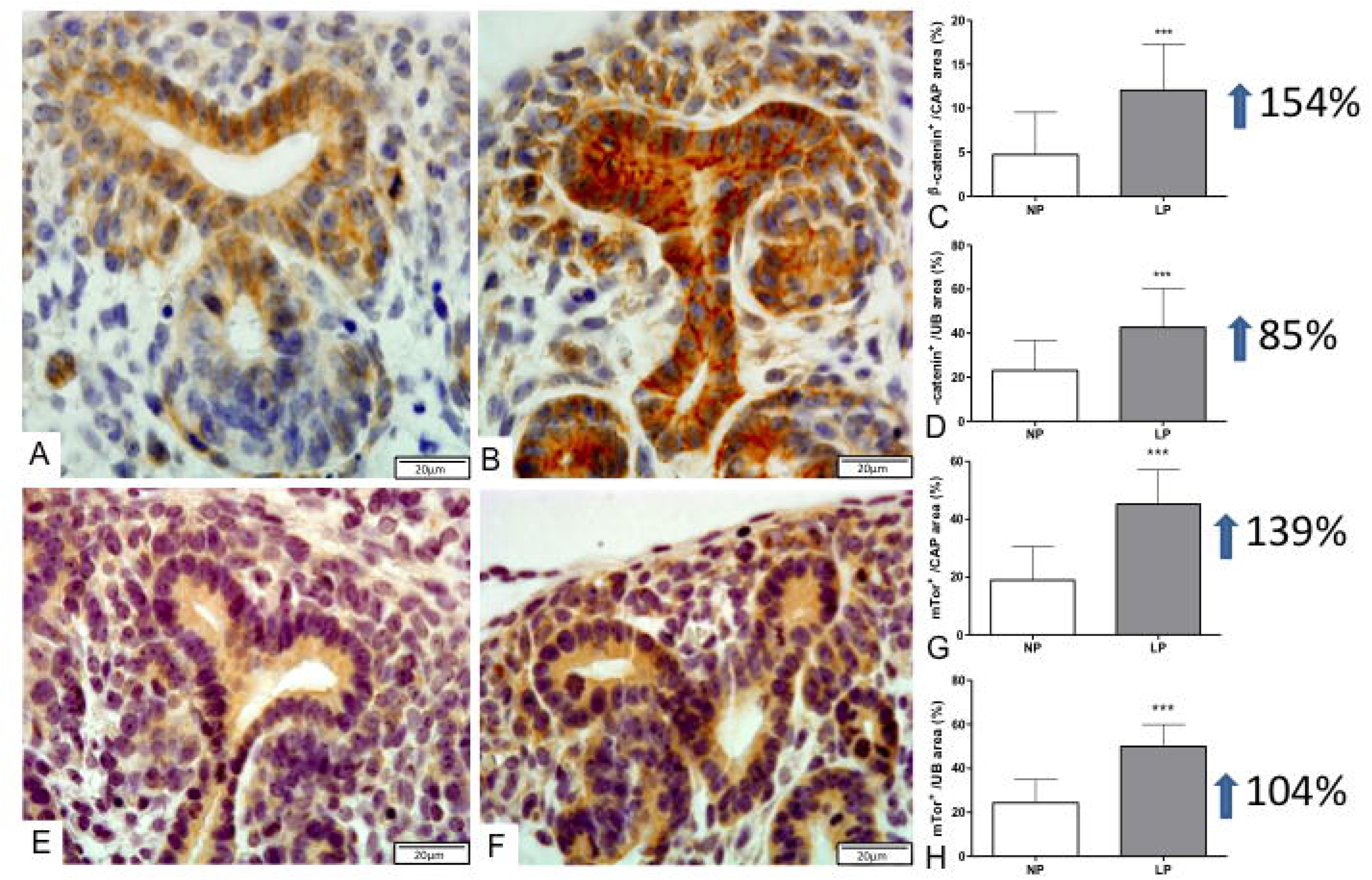
Immunoperoxidase for b-catenin and mTor in metanephros of 17DG fetus. In LP (B), the b-catenin labeled area was significantly raised in both CAP cells and UB epithelia (C, D) when compared to NP offspring (A). Also, mTor immunoreactivity occupied a more extensive area in LP (F) than in NP (E) offspring in analyzed structures (G, H). ***P<0.0001.

**Figure 8.**
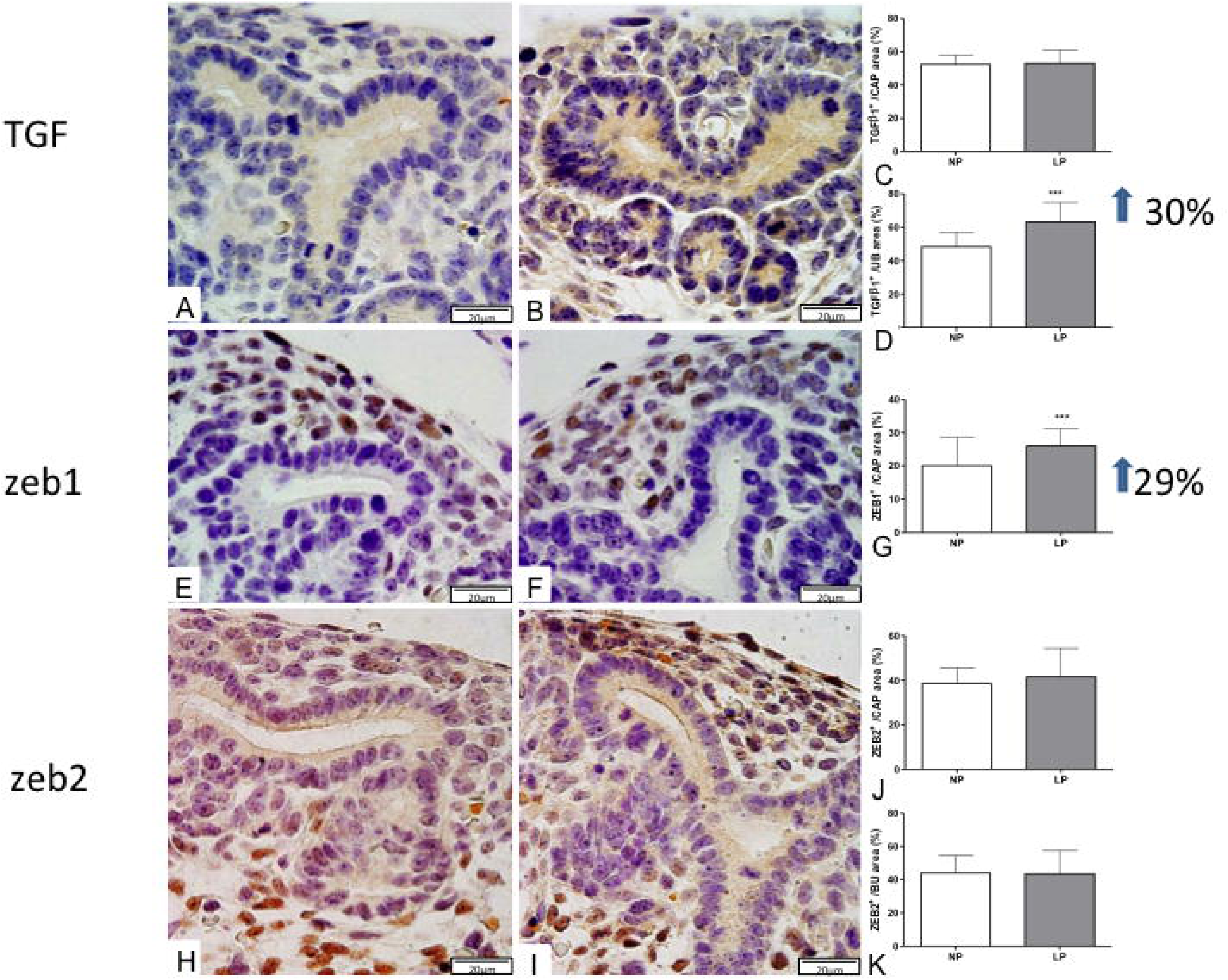
Immunoperoxidase for TGFb-1, ZEB1, and ZEB2 in metanephros of 17DG fetus. The area of TGFb-1 immunoreactivity in LP (B) compared to NP offspring (A) was not different in the CAP (C) but was significantly enhanced in UB (D). ZEB1 was detected in the nuclei of CAP, and other mesenchymal cells and, the LP (E) CAP occupied more extensive area (G) than in NP (F) offspring. The ZEB2 labeled area was not different hen compared to both groups in CAP (J) and UB (K). ***P<0.0001.

**Figure 9.**
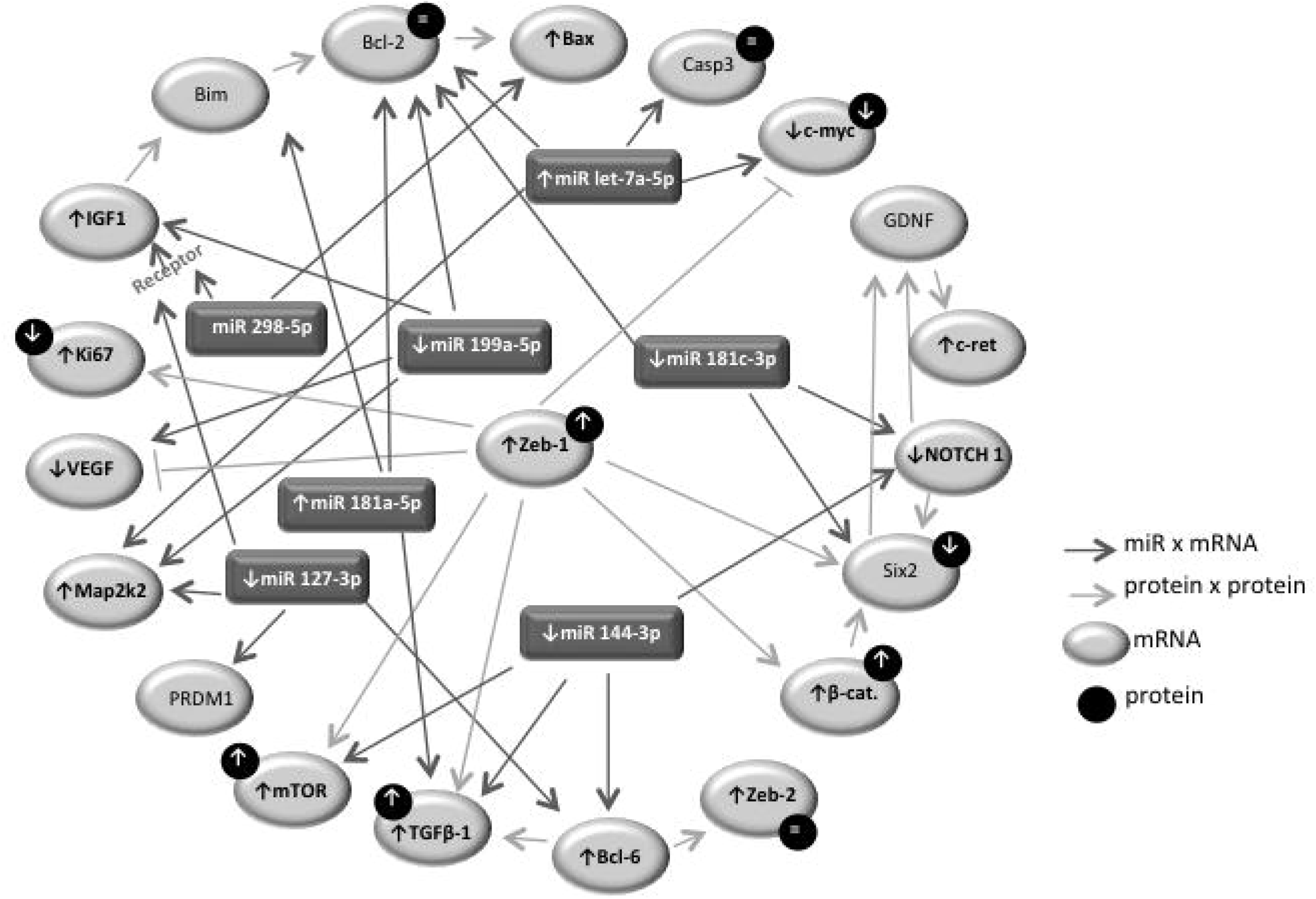
Deregulated miRNA-mRNA-protein pathways in metanephros of 17DG fetus from maternal restricted-protein intake.

## Discussion

The understanding of the cell and nephrogenesis molecular mechanisms has overgrown in late years (Cerqueira et al., 2017; Costantini and Kopan, 2010; Hendry et al., 2011; Kopan et al., 2014; Little and McMahon, 2012). However, many regulatory factors and signaling pathways involved in renal ontogenesis maintain unclear (Wang et al., 2018). It is known that miRNAs play a crucial role in regulating gene expression during the renal developmental processes (Lv et al., 2014; Marrone and Ho, 2014; Phua et al., 2015b; Wei et al., 2014). So far, to our knowledge, no prior study has performed an integrative miRNA and mRNA expression profiling analysis in maternal low-protein intake 17-DG male metanephros.

In the current study has proposed a novel molecular pathway that may be involved in the early nephrogenesis interruption and, consequently, reducing the number of nephrons. The present study, by NGS, identified 44 differentially expressed miRNAs, which 19 miRNA were up- and 25 downregulated in LP compared to NP offspring metanephros. Among the top 10 deregulated miRNAs, the study selected 7 miRNAs which its biological targets were related to proliferation, differentiation, and cellular apoptosis processes. As could be seen below, both miRNA-Seq and TaqMan data analysis have shown a consistent and specific change of miRNA expression in LP animals relative to control NP age-matched animals. The miR-181 family is composed of four highly conservative members, miR-181a, miR-181b, miR-181c, and miR-181d (Ji et al., 2010). In neoplastic cells, miR-181a plays the role of the tumor suppressor gene, inhibiting the proliferation and cell migration and, inducing cellular apoptosis (Chen. et al., 2010). In the current study, increased expression of miR-181a-5p in 17-DG LP relative to age-matched NP offspring; also, here, was demonstrated a 2-fold enhanced Bax/Bcl2 mRNA ratio in LP compared to NP offspring. These findings reaffirm increased CM apoptosis activity, although caspase mRNA expression is not altered, which may indicate a post-transcriptional mechanism on apoptosis regulation. Previous studies have shown that the BCL family promotes cytochrome release from mitochondria, and then inhibit the activation of Casp3 and subsequent cellular apoptosis (Scorrano and Korsmeyer, 2003). Li and colleagues, in an acute lung injury model, have shown that overexpressed miR-181a is related to decreased Bcl2 protein level; conversely, the inhibition of miR-181a increase Bcl2 levels (Li et al., 2016). The current study confirming Lv et al. (2014) demonstrated that miR-181c regulates negatively, the expression of Six2 expression and cell proliferation, in parallel to the loss of mesenchymal cells phenotype during kidney development in LP 17-DG offspring.

Studies from Xiang et al. demonstrate that miR-144 expression was decreased in kidney carcinoma cells (Xiang et al., 2016). On the other hand, they also showed that enhanced miR-144 expression suppresses renal carcinoma proliferation and decreasing the G2/M phase cells ratio. Our results show reduced expression of miR-144-3p in 17-DG LP offspring CAP, suggesting that it may be inducing cell proliferation.

Additionally, in renal cancer, the mTOR pathway plays an important pathogenic role in proliferation and response apoptosis during tumor development (Bailey et al., 2014; Ribback et al., 2015). Moreover, another study was demonstrated that the mTOR signaling pathway is crucial for the decreased number of nephrons in fetal?????, fetal kidneys whose mothers were subjected to the nutrient restriction (Nijland et al., 2007). The Xiang group demonstrated a significant molecular correlation between miR-144 and mTOR. They observed that overexpression of miR-144 inhibits the mTOR gene and protein expression. Conversely, the lower expression of miR-144 enhances mTOR mRNA expression (Xiang et al., 2016). Here, LP animals demonstrated an increased expression of mTOR. This finding suggests that the miR-144 expression may regulate mTOR, but not only by it, to promote cell proliferation.

The mammalian target of rapamycin complex 1 (mTORC1) is known to be indispensable for healthy embryonic development, has how to function to regulate the balance between growth and autophagy in physiological conditions and environmental stress (Gürke et al., 2016). It has been suggested that mTOR signaling plays a central role in the perception and response to intracellular nutrient availability (Xiang et al., 2016). Thus, this pathway would undoubtedly be involved in cellular responses in maternal protein underfeeding animals. The previous study demonstrated that an increase in mTOR was fundamental in reducing the number of nephrons in fetal kidneys, whose mothers were subjected to a nutrient restriction (Nijland et al., 2007). However, Volovelsky et al. (2018) reported that conditional deletion of mTOR in nephron progenitor cells profoundly disrupted nephrogenesis, and hemizygous deletion led to a significant reduction in nephron endowment. In the present study, the mRNA expression for mTOR in 17-DG LP offspring was remarkably increased, and the protein immunoreactivity was also enhanced, respectively, 139% and 104% in CM cells and UB.

The miR-127 has an essential role in proliferation, differentiation, and development, and one of its targets the proto-oncogene Bcl6 (Chen et al., 2013). Pan et al., show that in liver cells, that miR-127 underexpression is related to increased cell proliferation, and its overexpression was associated with proliferation inhibition (Pan et al., 2012). Also, Chen and col. classified this miRNA as a new regulator of cell senescence via Bcl6 (Chen et al., 2013). The reduction of miR 127-3p in 17-DG LP offspring was accompanied by increased mRNA for Ki67 associated with an increase of Bcl-6 in CM.

In this study, we also observed that the animals from the LP group presented a reduction in the nephrogenic area, corroborating the findings of Menendez-Castro and col. (2013 and 2014), who observed that both proliferative activity and the nephrogenic zone were significantly reduced in animals that suffered protein restriction (8.4% casein) relative to the controls group. We also observed a significant reduction of cells positively labeled for Ki67 in CM in protein-restricted animals.

The study showed that overexpression of miR-199a-5p reduces cystic cell proliferation and induces apoptosis, in addition to controlling the cell cycle (Sun et al., 2015). Expression of miR-199a-5p is reduced in LP, and the transcription of Ki67 and Map2k2 are increased. These findings indicate that gestational undernutrition promotes differentiation in detriment of proliferation associated with increased Zeb2 mRNA expression. Zeb2 is a transcription factor that regulates the renal tubule morphogenesis acting on proximal tubule development and glomerular-tubular junction (Rasouly et al., 2017).

The let-7 miRNA family expression has been extensively studied in several fetal tissues and, priority is related with reduced proliferation and induced cell differentiation (Ambros, 2012; Bao et al., 2013; Copley and Eaves, 2013; Meza-Sosa et al., 2014; Nagalakshmi et al., 2015). It has been shown in higher organisms, enhanced let-7 levels during embryogenesis (Schulman et al., 2005), and also that let-7a mature form is upregulated during the developmental mouse brain (Wulczyn et al., 2007). A study from Nagalakshmi et al. (2015) has shown that let-7 miRNAs potentially upregulated UB epithelial cells fate from precursor to the differentiated state. On the other hand, Yermalovich et al. (2019) have demonstrated the overexpression of Lin28b, an RNA-binding protein, is associated with suppressive let-7 miRNAs expression and elongated nephrogenesis, via the let-7 miRNAs upregulation. Thus, in the current study, the enhanced let-7a-5p miRNA expression in LP offspring might be associated with reducing CM cell proliferation, compromising the whole nephrogenesis relative to the NP group. While the heterochronic genes lin28 and let-7 are well-established regulators of developmental timing in invertebrates, their role in mammalian organogenesis was not fully understood. In the present study, we unprecedently report the reciprocal Lin28b/let-7 axis relationship during fetus kidney development in LP rats. So, we may hypothesize that overexpressed let-7 miRNAs, direct or indirectly, throughout a transient reduced LIN28B, might decrease the nephrogenesis process, and consequently, the kidney functional unit numbers potentially via upregulation of the Igf2/H19 locus. Conversely, a supposed kidney-specific regular expression of Lin28b normalizes whole kidney development, as was suggested in this study.

Here, such as in previous studies, was also observed increased let-7a-5p miRNA in parallel to the fall of c-Myc expression (Chang et al., 2008; Sampson et al., 2007). Prior study has demonstrated the involvement of MYC proteins in proliferation, growth, apoptosis and cell differentiation, critical events during renal organogenesis (Couillard and Trudel, 2009). In the LP 17-DG offspring, metanephric c-myc gene expression was reduced, and the CM c-myc immunoreactivity area was either 14% smaller, both when compared to NP offspring. Coincidently, CM cell number was also 14% reduced in LP, and KI-67 immunoreactivity was decreased even (48%) in LP relative to the NP group. Sustaining the present findings, authors have shown that c-myc plays a role in the developmental process, being crucial in the final phase of the UB branching by modulation of CM progenitor cell proliferation (Couillard and Trudel, 2009). It seems that let-7 should be strongly expressed at specific late stages of differentiation, but keeps downregulated in stem cells maintaining these cells in an undifferentiated state. Thus, the current study established that the let-7 family of miRNAs promotes MYC expression by a transcriptionally induced let-7 repressor, LIN28 enhancement and, post-transcriptional expressed LIN28 RNA binding-protein, promoting downregulated upon LP kidney cells differentiation (Figure 10).

**Figure 10.**
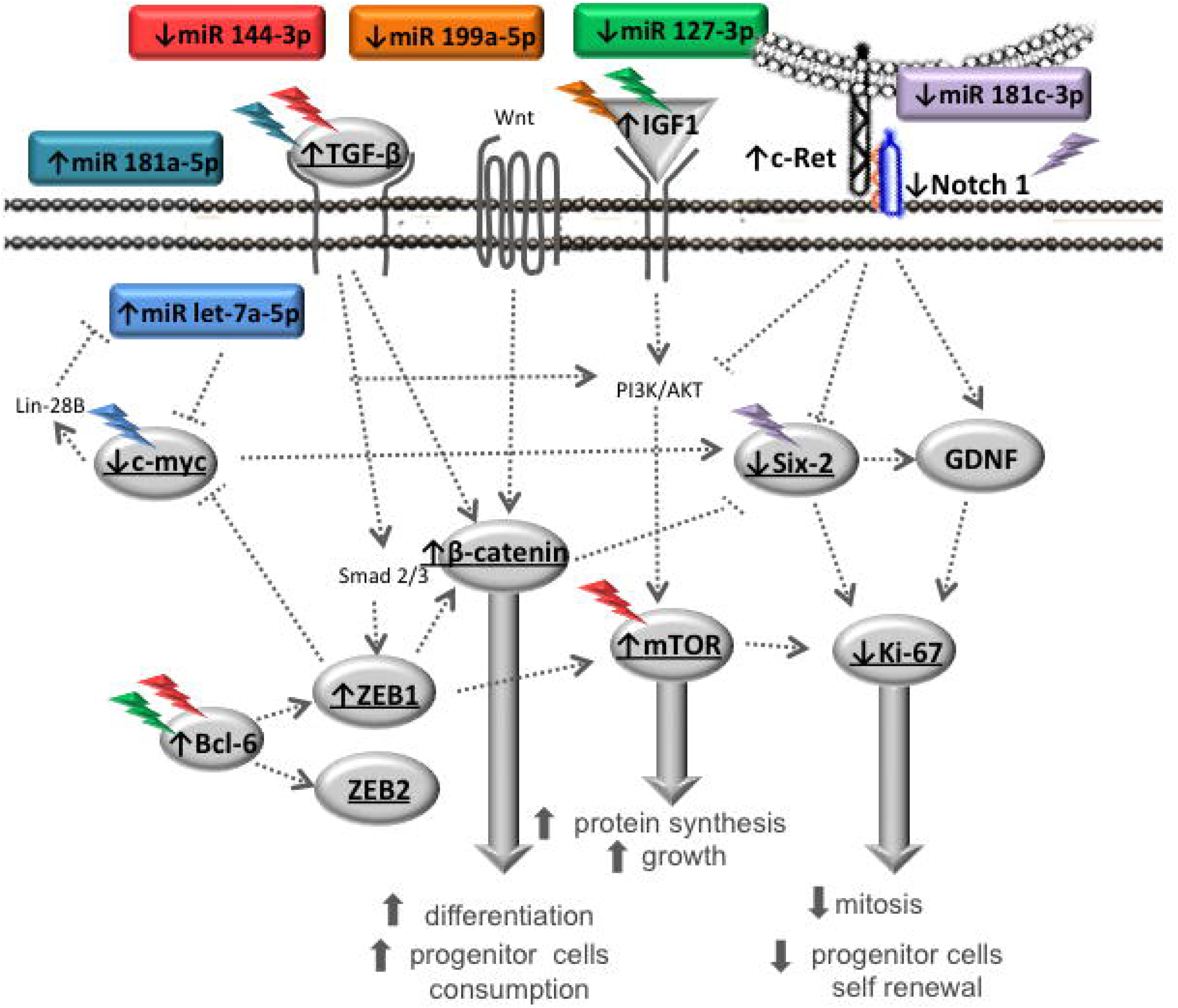
The picture depicted a schematic representation of deregulated pathways and biological response in metanephros of 17DG fetus from maternal restricted-protein intake.

The previous study has demonstrated that Six2 gene expression is downregulated during fetal ontogenesis when associated with reduced numbers of nephrons, arterial hypertension, and chronic renal failure in adulthood (Fogelgren et al., 2009). In parallel, the present study shows a significant 28% reduced Six2 positive cells in 17-DG LP compared to NP offspring, in similar proportion to previously observed decreased nephrons numbers in restricted-protein offspring. Six2, a specific marker of the renal stem cell system, is required to maintain under control the fate cell and the nephron progenitor cells population by suppressing signals induced-epithelial differentiation (Herzlinger et al., 1992). Its expression maintains a pool of CM cells available in the undifferentiated stage during renal development (Oliver et al., 1995; Self et al., 2006; Humphreys et al., 2008). The present data is sustained by a prior study in c-myc transgenic mice kidneys (Couillard and Trudel, 2009), that showed a decreased c-myc and six-2 immunopositive CM cells associated with reduced CM stem cell proliferation in 17-DG LP offspring.

Notch signaling is involved in the negative regulation of Six2, suggesting profound Notch impact on the gene regulatory network on the maintenance of nephron progenitor cells (Cheng, 2003; Cheng et al., 2009). Once that reduced expression of Six2 is required for differentiation of the nephron progenitors, the Notch signaling is fundamental to downregulate Six2. Thus, we may argue that reduced c-myc and Notch signaling in this study might stimulate renal cell differentiation since it regulates the size of the CM cell population. In several cases, the Wnt/β-catenin pathway initiates the differentiation of nephron progenitor cells being, however, essential for the maintenance of nephron progenitor cells (Brown et al., 2015). Thus, a low level of β-catenin might be required for the maintenance of Six2 expression, and a high level of β-catenin may contribute to the downregulation of Six2. Wnt/β-catenin and Notch signals may coordinate that Six2 expression regulation.

Additionally, elevated levels of β-catenin activity also determine nephron progenitor cell fate. Usually, only when a sufficient pool of CM cells is reached, nephrogenesis proceed. In this way, we assume that 17-DG LP offspring has a 28% reduction of CM, which will affect the percentage of nephrons in adulthood. Thus, partial conclusions of the present study sustain that, in LP offspring metanephros, the increased let-7a-5p and β-catenin expression and reduced Notch signal modulate the c-myc, six2, and KI-67 leading to reduction of progenitor cells self-renewal. By the way, the exhaustion of the remaining CM progenitor cell endowment, in turn, predisposes to reduced nephron numbers, arterial hypertension, and renal disorders in adult age (Figure 10).

As previously observed in our Lab, a reduction of 28.3% in ureteric bud branches after 14.5 days of gestational protein restriction (Mesquita et al., 2010). Genes encoding the peptide growth factor glial-derived neurotrophic factor (GDNF), c-Ret tyrosine kinase receptor, and its coreceptor Gfra1 are recognized as crucial regulators of ureteric bud outgrowth (Davis et al., 2015). Thus, the c-ret receptor, when activated by GDNF, induces UB branching. In the 17-DG offspring, was observed a significant increase in c-Ret receptor coding mRNA, which would theoretically lead to a rise in UB ramifications. However, in the present study, the GDNF expression was unchanged, being plausible to suppose that, despite the increase of c-Ret, the UB branching did not also change. How is it known that Six2 regulates transcription of GDNF (Brodbeck et al., 2004), thus, the reduction of 28% in the cells positive for Six2 could affect, in the same proportion the GDNF expression which in turn, would act in the decrease of 28.3% of the ramifications of the ureteric bud as previously observed (Mesquita et al., 2010).

Karner et al. (2011) demonstrated that the signaling pathway is Wnt9b/ β-catenin is required both for nephron progenitor cell renewal and for differentiation, thus being essential for the formation of nephrons during embryogenesis. However, it is not yet known how β-catenin regulates these two processes. This signaling pathway is evolutionarily conserved and plays an essential role in the development of organs, tissues, and injury repair in multicellular organisms. Although this pathway plays a crucial role in the development of the kidney, it is silenced in the kidneys of normal adults and is reactivated against some kidney damage (Halt and Vainio, 2014). c-myc is a transcriptional target of β-catenin, which in turn regulates the proliferation and differentiation of renal tubular epithelium (Hu and Rosenblum, 2005). Recently the Pan group showed that myc cooperates with β-catenin to enhance the renewal of nephron progenitor cells (Pan et al., 2017), our LP animals showed lower c-myc expression, both genetic and protein so we can assume that these animals had a lower reserve of renewing cells, which are necessary for proliferation and survival, and may reflect on the smaller number of nephrons that are seen in the model.

β-catenin also has been shown to participate in a signaling pathway that controls organ size. This protein modulates the CM progenitor cell population and, together with a transcriptional complex, binds to the promoter region of the bcl2l1 anti-apoptotic gene in the developing kidney (Boivin et al., 2015). Besides, β-catenin interacts with the c-ret receptor, which, when stimulated by GDNF, releases it into the cytosol, and β-catenin migrates to the nucleus of UB cells (Sarin et al., 2014). In addition to its participation in the Wolffian duct elongation, β-catenin is essential for UB branching and differentiation (Bridgewater et al., 2008; Marose et al., 2009). However, have been shown that the increase in β-catenin expression has led to increased TGFB1 expression in epithelial cells inhibiting UB branching and causing premature differentiation of CM progenitor cells (Bridgewater et al., 2011) Several studies have shown that increased β-catenin expression in CM disrupts UB ramifications and nephrogenesis by deregulating essential genes during renal development (Boivin et al., 2015). In the present study, the gene and protein expression of β-catenin is increased during the studied periods of renal development in LP animals. During renal development, β-catenin is expressed in both UB and CM modulating the gene programming that governs new budding and UB branching. In 17-DG LP is evident the increase of beta-catenin in the UB in the cytosol, whereas, in NP, the location is in the nucleus. Here, in CM, we also observed β-catenin labeling but not in NP age-matched offspring.

Although several authors have widely studied the process of normal nephrogenesis (Constantini and Kopan, 2010; Little and Mcmahon, 2012; Mugford et al., 2010), little is known about the mechanisms that determine the number of nephrons. Moreover, how the balance between nephron progenitor renewal and differentiation is essential for the proper development and function of the kidneys, this is a particularly important issue, because failure to achieve adequate numbers of nephrons is a risk factor for chronic renal disorder (Benz et al., 2011; Hoy et al., 2006; Luyckx and Brenner, 2005; Pan et al., 2017). In conclusion, the present study demonstrates that the miRNAs, mRNAs, and proteins are modified in LP animals with 17 DGs leading to the reduction of the reciprocal induction between CM and UB, and hence the number of nephrons (Figure 11). These findings will facilitate new functional approaches and further studies to elucidate the regulatory mechanisms involved in the processes of proliferation, differentiation, and apoptosis that occurs during renal development.

**Figure 11.**
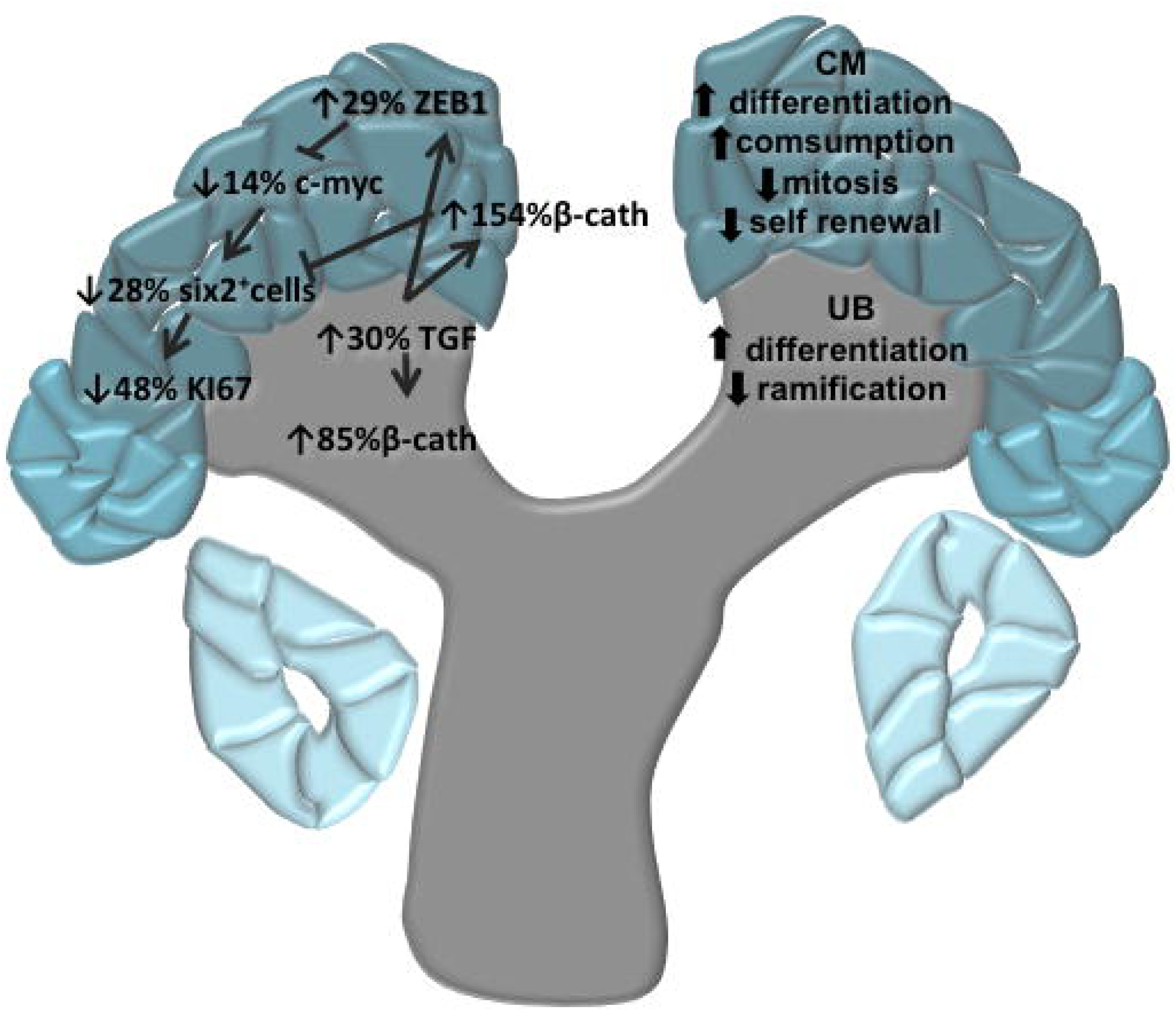
The picture depicted a schematic representation of supposed factors that evolved in a 28% reduction of the CM stem cells and nephron number in maternal restricted protein intake.

## Acknowledgments

We thank the access to equipment and assistance provided by the National Institute of Science and Technology on Photonics Applied to Cell Biology (INFABIC) at the State University of Campinas; Data analysis was partial and generously performed in collaboration with Tao Chen, Ph.D. from the Division of Genetic and Molecular Toxicological, National Center for Toxicological Research, Jefferson, AR, USA. This work was supported by Fundação de Amparo à Pesquisa do Estado de São Paulo (FAPESP, 2013/12486-5 and 2014/50938-8), Coordenação de Aperfeiçoamento de Pessoal de Nível Superior (CAPES) and Conselho Nacional de Desenvolvimento Científico e Tecnológico (CNPq, 465699/2014-6).

